# Tuberculosis susceptibility and inbreeding depression hinder *ex-situ* conservation in a critically endangered rainforest bird

**DOI:** 10.1101/2025.08.17.670676

**Authors:** Peri E. Bolton, Dustin J. Foote, Nancy E. Drilling, Susan B. McRae, Michael D. Sorenson, Christopher N. Balakrishnan

**Author notes:** these authors contributed equally to this work (Peri E. Bolton, Dustin J. Foote).

## Abstract

Captive breeding can be a key component of species conservation strategies, but also exposes these rare species to novel environments including the pathogen landscape. The critically endangered white-winged wood duck (WWWD) *Asarcornis scutulata* has experienced substantial population declines, local extirpations and fragmentation of its former range in South-East Asia, making it one of the rarest birds in the world. Like other rare species, WWWD declines have led to the initiation of captive breeding programs, but these have been hampered by the WWWDs’ high susceptibility to *Mycobacterium avium*, avian tuberculosis (TB). In this study we describe genome-wide patterns of diversity to understand the WWWD’s demographic and phylogeographic history, inbreeding in the wild and in captivity, and the causes of TB susceptibility. Captive birds, which originated from northeast India, are genetically differentiated from wild birds sampled in Sumatra, Indonesia, likely reflecting long-standing phylogeographic structure. Demographic analyses revealed that long-term (Pleistocene) population declines preceded anthropogenic declines, a pattern shared with other codistributed, forest-dependent species. All sampled WWWD populations had extremely low genetic diversity but wild-sampled birds retained higher Major Histocompatibility Complex (MHC) diversity, reflecting important functional diversity in the wild. Genetic diversity has eroded over time in captivity and importantly, birds with higher levels of inbreeding succumb earlier to TB infections, suggesting inbreeding depression. Finally, by comparing gene expression between susceptible WWWD and resistant redhead ducks *Aythya americana* we identify possible mechanisms of TB susceptibility in WWWD. Altogether our study provides genomically-guided objectives for future management and a cautionary tale for *ex-situ* conservation.

## INTRODUCTION

Captive breeding is an important component of species conservation strategies. However, husbandry in captivity poses additional challenges as these rare species are exposed to novel environments, including pathogens (1). These novel exposures may interact with inbreeding to hinder captive breeding success. Conservation genomics has the potential to provide not only an understanding of the demographic history of rare and threatened species but also their ability to adapt and respond to novel threats in a changing environment (2). However, genomic efforts have mostly focussed on species in the global north (3), leaving many equally or more imperiled species in the tropics poorly understood.

*Asarcornis scutulata*, the white-winged wood duck (Figure 1A), is one of the rarest birds on earth, yet we know nothing about patterns of genetic variation in this species. Also known simply as the white-winged duck, the longer moniker is more commonly used within the species’ range (4) so we use that nomenclature here, abbreviated as WWWD. The current WWWD range extends from Northeast India to Sumatra, though only small, fragmented populations persist across this range (Figure 1B) (4–7). Somewhat unusual for waterfowl, WWWDs are forest dwelling and cavity nesting. WWWDs are highly dependent on mature rainforest with slow moving water bodies, habitats that have been particularly degraded over the last century. Deforestation and poaching remain the biggest threats to the survival of the species.

**Figure 1.**
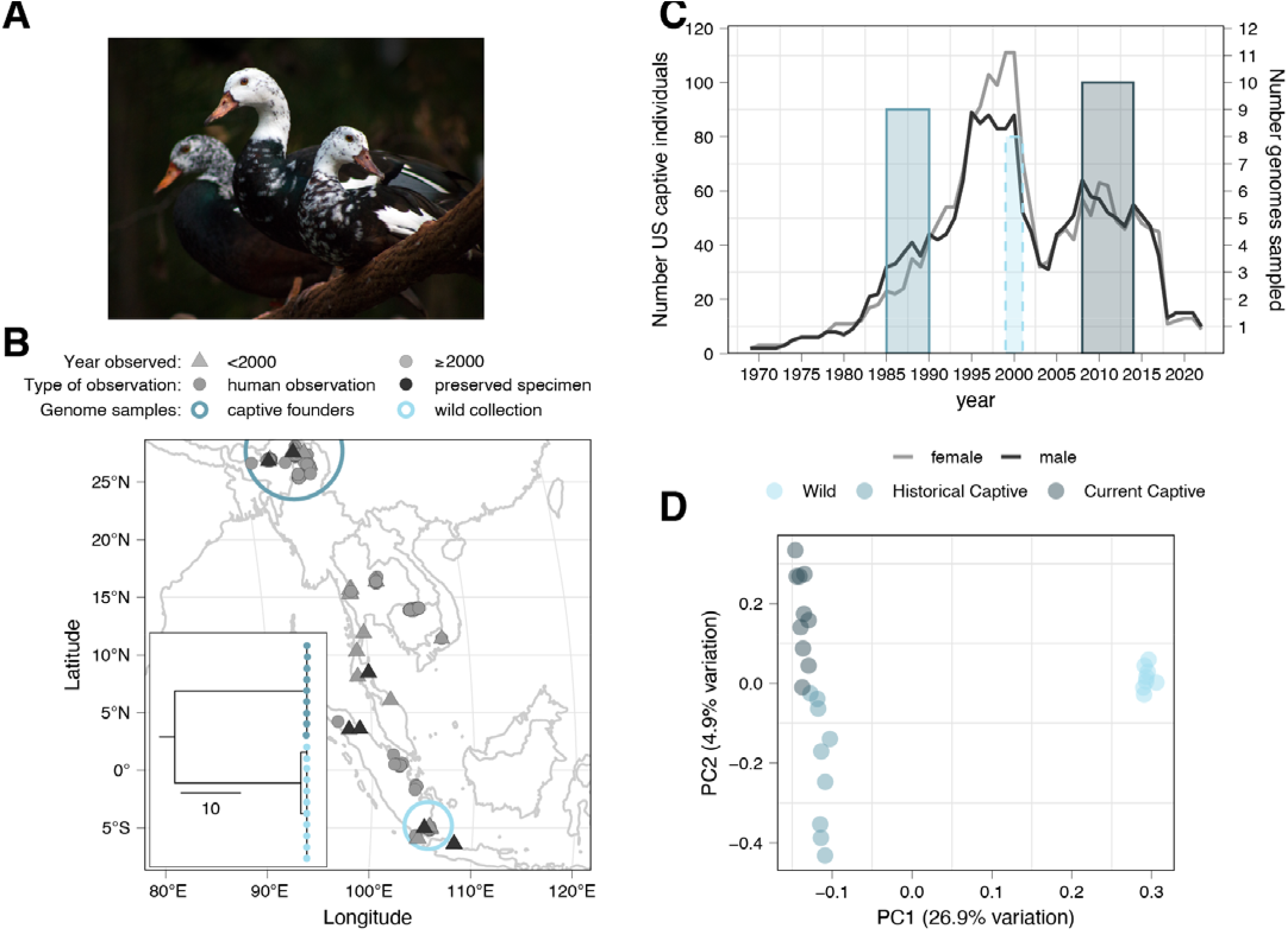
Endangered white-winged wood ducks have experienced population declines in the US captive breeding program, and are genetically differentiated from wild ducks. A) White-winged wood ducks (*Asarcornis scutulata*) at Sylvan Heights Bird Park, North Carolina. Photo: Katie Lubbock. B) Time series showing US captive population size of male (black lines) and female (grey lines) white-winged wood ducks (left y-axis). Colored rectangles indicate the hatch years for captive birds, and sampling period for wild birds for the genetic samples used in this study (x-axis) and their height indicates the number of samples used in this study (right y-axis). Three additional wild birds were included in mtDNA phylogeny. C) Distribution of white-winged wood ducks in south-east Asia from Global Biodiversity Information Facility (GBIF) records (79). Black points indicate museum specimens, whereas grey indicate human observations such as from eBird. Triangles show observations prior to the year 2000 indicating likely extinction in Java and Malay Peninsula. Observations in central northern Thailand are from a reintroduction program [7]. Large colored circles indicate the approximate locations of the founders of the captive population in Assam (dark blue), and the feather samples taken from wild ducks in Sumatra (light blue). Inset: Maximum parsimony mitochondrial DNA phylogeny of wild and captive birds, where scale is the number of changes along each branch. D) Principal components analysis of autosomal variants for individuals used in this study, color coded by population.

Recent estimates suggest only 150 to 450 mature adult WWWDs remain in the wild, with the largest populations in northern Myanmar and Arunachal Pradesh and Assam Provinces in India (8). The WWWD is now considered critically endangered by the International Union for Conservation of Nature (IUCN), but protection efforts in India began in 1937 and the species was placed on the Indian Special Protected List in 1952 (9). The World Wildlife Fund selected the WWWD for a focal species project in 1968. As a result, the International Waterfowl Research Bureau and the World Wildlife Fund recommended that immediate action was needed to ensure the survival of WWWD in Assam (6, 9, 10).

As part of a resulting conservation plan, captive populations were initiated in the United Kingdom, and then the United States. These captive populations were established using birds collected in Assam in 1969 and in 1970 (9). Both of these early wild collections took place on the same tea plantation in upper Assam (Figure 1B). Staff at two Wildfowl and Wetland Trust (WWT) centers, located at Slimbridge and Peakirk, UK, were able to rapidly grow their captive WWWD populations to a total of 86 individuals by 1976 (11). Soon after establishment of the captive breeding program, however, it became clear that *Mycobacterium avium* — avian tuberculosis (TB) — was the source of mortality for most individuals. Between 1976 and 1991, 102 out of 121 birds (84%) succumbed to TB (12). Originating from the same wild population in Assam, WWWDs were initially imported to various facilities in the US. Led by Sylvan Heights Bird Park (SHBP) in Scotland Neck, North Carolina, the US population expanded to just under 200 individuals (Figure 1C) but following this population growth (*circa* 1999), avian TB began to significantly affect the North American population, primarily due to an increase in mortality in birds aged 2-6 years (13). This rapid turn of fortune for the US captive population suggests the possibility that inbreeding depression may have interacted with TB exposure to reduce the viability of captive breeding. At present, the current captive population is descended from just two wild founders.

In this study, we set out to describe genetic variation in WWWDs, including wild birds sampled in Indonesia and captive birds in the US, originally sourced from India. We aimed to provide a first look at the extant genetic diversity in this species with an aim to guide future conservation efforts. We specifically explore genetic variation in immune system genes of the major histocompatibility complex (MHC) and transcriptome-wide expression patterns to provide mechanistic insights into TB susceptibility in WWWDs.

## RESULTS

### Whole genome sequencing

We generated whole genome sequencing data for samples from three populations: “Historical Captive” (n = 9) birds were hatched at multiple breeding facilities in the U.S. between 1985 and 1990, and were later housed together at SHBP. “Current Captive” (n = 10) birds hatched between 2008 and 2014 at SHBP (Figure 1B). “Wild” Indonesian (n = 11) samples were collected between 1999 and 2001 in the Way Kambas National Park in Lampung Province, Indonesia (southern tip of Sumatra, Figure 1C; Table S1). We used two different reference genomes and read mapping strategies for different analyses. For analyses of mitochondrial DNA (mtDNA) and pairwise sequentially markovian coalescent (PSMC) analyses we used the draft quality WWWD genome assembly, which includes a complete mtDNA genome (GCA_013398475.1 (14), see methods). For other population genomic analyses, we used a chromosome-level tufted duck (*Aythya fuligula*) genome assembly (GCF_009819795.1 (15)) to allow insights into complex genomic regions like the major histocompatibility complex (MHC). Overall mapping rates were 97.63% to the *A. fuligula* assembly, marginally higher than mapping to the less contiguous WWWD assembly. Captive birds were sequenced to an average of 10.5x depth and wild birds were sequenced to an average 16.5x coverage; we downsampled to equal read-depths as needed for specific analyses (see methods). All resequencing data have been deposited in the NCBI Short Read Archive under PRJNA1258233. Three wild samples were removed from genome-wide analyses due to poor data quality (mapping rates < 50%).

### Indonesian and Indian birds are genetically distinctive

Analyses of both mitochondrial DNA (mtDNA) and genome-wide variation indicate clear, historical divergence of the Indian-derived, captive population and the wild Indonesian population. Mitochondrial lineages are reciprocally monophyletic, with 46 to 48 differences (∼0.28 % divergence; Figure 1C) between the captive and wild lineages. A time-calibrated phylogeny (Figure S1) suggests that these lineages diverged during the late Pleistocene at ∼170,000 years ago (95% HPD range = 103-241 kya). We note that this estimate is considerably more recent than the calibration point, such that it likely overestimates the actual divergence time to some degree (16). Similarly, Principal Components Analysis of genome-wide variants using PCAngsd clearly separated the three sampled populations (Figure 1D). Captive (India-derived) and wild (Indonesian) birds separated along PC1 (26.9% of variation), whereas the historical and current samples of captive birds separated along PC2 (4.9% of the variation) (17). This separation in PC space corresponds to moderately high autosomal FST values between captive and wild birds (historical versus wild = 0.3, current versus wild = 0.4) and relatively low genome-wide differentiation between current and historical captive birds (0.04).

### Genetic diversity is low and inbreeding is high

Demographic analyses using *PSMC* (18) suggest long-term population declines, particularly for the sample of wild Indonesian birds, which show evidence of steady population decline throughout the Pleistocene (Figure 2A). In contrast, the WWWD population in northeast India may have maintained a larger population size during the late Pleistocene. Interestingly, the point at which the demographic trajectories of these two populations diverge, around 150,000 years ago, may correspond to the isolation of these populations and their mtDNA lineage divergence. Effective population sizes for the Indian population remained relatively large until about 100,000 years ago after which this population also declined.

**Figure 2.**
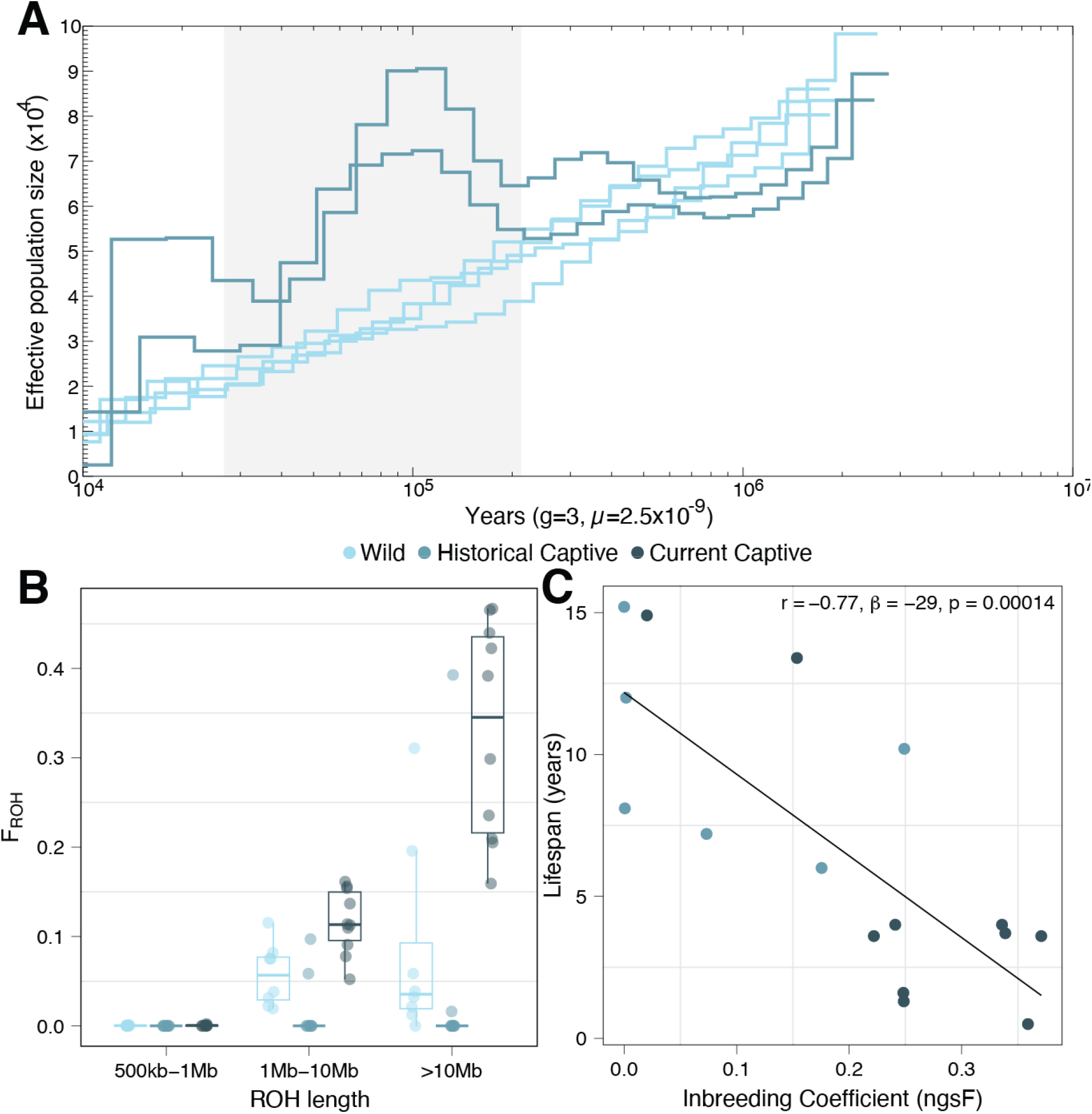
Populations of white-winged wood ducks from India and Indonesia have declined during the Pleistocene, and have experienced ongoing inbreeding and inbreeding depression. A) Estimates of effective population size through time from PSMC for wild (Indonesia) and historical (India) captive ducks. Grey shaded box shows the late Pleistocene period (11.7-115kya), with years scaled by generation time of three years and a per-generation mutation rate of 2.5 × 10^-9^ (55). B) Inbreeding coefficients calculated from runs of homozygosity (ROH) in genomic size-classes per individual. C) Inbred birds in captivity typically live shorter lives (see also Table S2 and Figure S4-S6).

Consistent with both longer term and more recent, anthropogenically-driven declines, nucleotide diversity (π) was very low in all three population samples, with current captive birds (median π = 0.0008) harboring about 53% of the diversity of their captive ancestors (median π = 0.0015). Surprisingly, genetic diversity in the wild-sampled birds was similarly low (median π = 0.0010) (Figure S2). Autosomal heterozygosity estimates showed similar trends, with the wild population having lower heterozygosity than the historical captive population and the current captive population declining considerably in heterozygosity (∼50%) (wild het = 0.001, historical het = 0.002, current het = 0.001; Figure S2). Consistent with patterns of nuclear diversity, a single mtDNA haplotype is found in the captive population, whereas the wild sample included two closely related haplotypes (Figure 1B). These patterns are also reflected in the lengths and abundance of runs of homozygosity (ROHs), which reflect inbreeding (19). Recent inbreeding produces longer segments, which may be broken into shorter segments by subsequent recombination. Current captive birds had the largest number of long (>10Mb) segments and highest inbreeding coefficients as measured by F_ROH_ (F_ROH_ = ΣL_ROH_/L_autosomes_ = 0.47 median for the current captive population), whereas ROHs were observed in only two historical captive individuals (median F_ROH_ = 0). Some of the wild WWWDs had many long ROHs, indicating recent inbreeding, but with a lower level of inbreeding overall (median F_ROH_ = 0.11). This is in broad agreement with individual inbreeding coefficients from *ngsF* (Figure S3), which were very low in the historical captive sample (median F = 0.007), and much higher in contemporary captives (median F = 0.25). In contrast, wild birds had the highest individual inbreeding coefficients (median F = 0.36), primarily due to intermediate length ROHs, which may indicate ongoing inbreeding in a smaller and declining population on the island of Sumatra.

For a sample of sixteen captive birds with known age at death (ten current and six historical captives), we observed a strong and significant negative relationship between the *ngsF* individual inbreeding coefficient and age at death (Figure 2C; r = - 0.8, β = -29, p = 0.00014, see also F_ROH_, Figure S5, Table S2). More highly inbred individuals died earlier, usually from mycobacteriosis. This pattern is driven in large part by a difference between the historical and recent captives, with the recent captives dying younger than their historical counterparts. Notably, the two recent captives with the longest lifespans were the least inbred.

### MHC diversity is low in captivity

Given the effect of inbreeding on susceptibility to mycobacteriosis, we next examined genetic diversity in the major histocompatibility complex, a part of the genome with a key role in immunity broadly and TB specifically (20, 21). Due to MHC copy number variation among bird species we estimated diversity with two approaches: 1) using reads aligned to the heterospecific tufted duck reference genome, which may underestimate diversity due to reads that fail to align, and 2) using sample-depth corrected MHC reads aligned to a single MHC copy (see methods).

Using the latter dataset, we estimated MHC gene copy numbers for WWWDs. Coverage of reads mapped to a single MHC copy was different among populations, especially for MHC Class I (Table S3). Based on these differences in relative sequencing depth, we estimate that wild Indonesian WWWDs have at least three MHC class I copies, and 1-2 class IIβ copies. In contrast, captive ducks have approximately half as many copies of class I (1-2), but a comparable number of class IIβ copies.

There was a significant loss of heterozygosity in the peptide binding region (exons 2-3) of MHC class I genes in captive birds (Figure 3A; Figure S7), with a non-significant loss in nucleotide diversity (Figure 3B; Figure S6). These patterns are similarly reflected in patterns of amino acid diversity (Figure 3C-D), suggesting a loss of functional diversity. MHC reads aligned to single gene copy suggest a reduction in MHC Class II PBR (exon 2) heterozygosity and nucleotide diversity (Figure 3A), but this was not evident when the data were aligned to the tufted duck genome (Figure S6), perhaps due to variation in read depth across paralogs (Figure S8). Importantly, wild birds have relatively high MHC diversity at both Class I and Class II loci.

**Figure 3.**
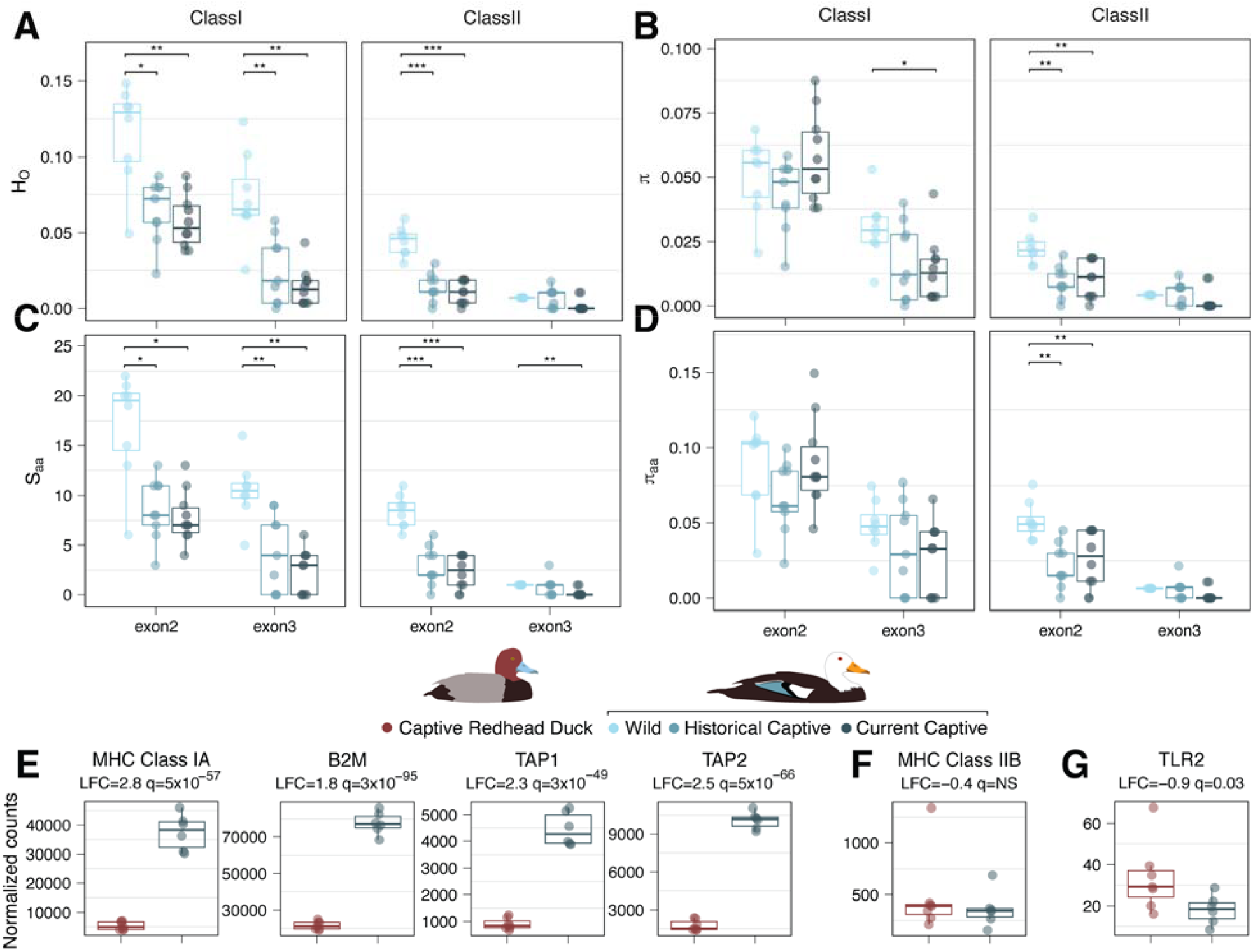
White-winged wood ducks in captivity have lost nucleotide and amino acid diversity in Major Histocompatibility Complex genes, and show potential immune gene dysregulation. A-D) Diversity of peptide binding exons in the Major Histocompatibility Complex (MHC). Estimates based on aligning individual data to single exon sequences for each class. A) Per-site heterozygosity, B) Nucleotide diversity, C) Number of variable amino acids, D) Pairwise amino acid diversity. Reductions in MHC Class IIB diversity are less dramatic when aligned to the tufted duck genome (Figure S7). E) MHC Class I and other key genes in the Class I antigen presentation pathway are significantly upregulated in WWWD compared to non-susceptible RH ducks. F) MHC Class II is not differentially expressed between these species. G) Among other differentially expressed immune genes, toll-like receptor 2 is downregulated in WWWD.

### Immune gene expression in white-winged wood ducks

Birds in captive environments, including those at Sylvan Heights Bird Park, are frequently exposed to *M. avium* but many species are only rarely affected by infections (22). To provide a window into the mechanism of susceptibility, we conducted RNA-seq on whole blood from WWWDs and closely-related redhead (RH) ducks *Aythya americana*, which rarely become sick from TB exposure. We examined both species-level differences in expression in asymptomatic birds (WWWD = 6, RH = 7) and differences in symptomatic (WWWD=2, RH=1) versus asymptomatic birds within species (Table S4).

We found that 56% of the whole-blood transcriptome was differentially expressed between asymptomatic redhead and asymptomatic WWWDs (5,797 genes, q<0.05, Table S5). There were no GO Biological Processes or KEGG pathways that were significantly enriched among these genes (Table S6; Table S7). There were 341 candidate immune genes (see methods) differentially expressed between species, which was not a significant enrichment (p=0.9). These immune genes included 27 RIG-I-like receptor signalling genes (ko04622), and 51 genes in the Tuberculosis KEGG pathway (ko05152) (p>0.05, Table S7). Among the differentially expressed candidate immune genes were four key MHC genes, MHC Class I⍰ and beta-2 microglobulin (B2M), which dimerize to form the Class I receptor protein, and transporter associated with antigen processing 1 and 2 (TAP1 and TAP2), which together form the antigen presentation complex. All four of these genes are substantially upregulated in asymptomatic WWWDs relative to redhead ducks (LFC=1.8-2.8, q<5×10^-57^, Figure 3E). Within the TB pathway, there was no evidence for differential expression in MHC IIβ (Figure 3F), but toll-like receptor 2 (TLR2) was downregulated in WWWDs (Figure 3G).

Although limited in sample size, we also contrasted symptomatic and asymptomatic WWWDs. In total, 186 genes were differentially expressed after excluding genes with expression patterns affected by outliers (q*<0.05; Table S8). There was significant enrichment for genes in the GO Biological Process “Regulation of Apoptotic Process” (GO:0042981, q=0.04, 5 genes). There was also weaker, non-significant, enrichment of genes involved in “Immune Response” (GO:0006955, q=0.1, 5 genes; Table S9), *Yersinia* Infection (ko05135, 5 genes, p=0.03, q=0.8) and Chemokine Signalling Pathway (ko04062, 5 genes, p=0.04, q=0.8, Table S10). We found fewer disease-related DEGs in redhead ducks (34 genes q*<0.05) (Table S11), with significant enrichment only non-immune KEGG “Replication and Repair” (ko03030, 3 genes, Table S12-S13). We found seven immune genes had opposite expression direction in symptomatic individuals of the different species (Figure S11). No genes with significant differences in disease expression were shared between the two species.

## DISCUSSION

With biodiversity disappearing faster than it can be described, we provide here the first characterization of genetic diversity in a critically endangered bird species. The white-winged wood duck represents a monotypic genus *Asarcornis (23, 24)* that historically ranged widely from eastern India to Indonesia. Though subspecies are not currently recognized (25), Indonesian WWWDs have been considered a separate subspecies (*leucoptera*) by earlier authors (26, 27) based on an apparently greater extent of white on their head and mantle plumage as compared to Indian ducks (nominate) (28). We found that captive birds derived from northeast India were substantially genetically differentiated from those sampled in Sumatra (Figure 1). The level of genetic divergence between Indian and Indonesian birds, including reciprocal monophyly of mitochondrial lineages, is consistent with the recognition of distinct subspecies. Given our sparse geographic sampling, however, the phylogeographic distribution of these lineages and whether and where there is a clear break between them remains unresolved. Further work is needed to test for associations between genetic and phenotypic variation.

We assessed genetic variation in WWWD populations, including temporal samples of India-derived captive birds in the US and wild birds sampled in Indonesia. All three WWWD populations had extremely low genetic diversity with levels similar to other critically endangered species like the five kiwi lineages (*Apteryx* genus) (29), crested ibis *Nipponia nippon* (30), and the channel island fox *Urocyon littoralis* (31). Unexpectedly, a sample of wild Indonesian WWWD collected in 1999-2001 had similarly low genetic diversity to the captive populations sourced from India, which have been affected by both founder effects and many generations of inbreeding in captivity. Low genetic diversity in the wild appears to be a result of long-term population decline driven by environmental change during the Pleistocene (Figure 2), and perhaps compounded by founder effects during colonization of Sumatra and/or subsequent isolation following sea-level rise and insularisation (32). Therefore, the low observed genetic diversity in the wild is not solely driven by recent anthropogenic effects, but signatures of recent inbreeding are likely signals of recent fragmentation (33). Instead, environmental change during Pleistocene glacial cycles and the associated reduction of forest habitats (34) appear to have initiated the decline of WWWDs globally. This observed demographic trajectory of WWWDs is similar to other tropical vertebrates (35, 36), including the critically endangered Sumatran rhino, which shares a similar geographic distribution and habitat requirements (37). As a charismatic megafauna species, the Sumatran rhino is a “flagship” for conservation efforts that may also promote the persistence of WWWDs.

The current captive US population of WWWDs is derived from a single pair of founders from Assam, India. As a result, we observed that genetic diversity has eroded substantially over time and that more highly inbred birds had shorter lifespans (Figure 2C). This pattern is largely explained by differences in lifespan between individuals in our current and historical samples, and so could be explained in part by possible increases in TB exposure over time. Among our “current” sample, however, the two birds with the longest lifespans were the least inbred, suggesting that inbreeding depression, and not just TB exposure, contributes to reduced lifespans. This finding suggests that genetic improvements to the current captive stock will be an important part of future WWWD *ex-situ* management (38). Most WWWDs breed at ∼3 years of age, and early in the breeding program, birds lived long enough to lay multiple clutches of eggs. This is no longer the case (13).

Despite having low overall genetic diversity, wild Indonesian WWWDs had higher MHC diversity than the captive population (Figure 3A-D), representing a potentially important reservoir of functional genetic variation. Similar retention of MHC diversity has been seen in other bird species that have suffered population bottlenecks during island colonization (39, 40). The observed difference in MHC diversity between captive Indian and wild Indonesian birds is due to an apparent difference in copy number, with further reductions in per locus diversity through time. Due to our sampling, however, it remains unclear whether the difference in copy number is an evolved difference between Indian and Indonesian birds, or is due to the two US founders from the Indian population having an atypically small number of loci. Sampling of wild, Indian birds would resolve this uncertainty.

Given the low MHC diversity in captive birds, we sought further insights into the mechanism driving differences in susceptibility between WWWDs and other birds housed in the same conditions. By conducting RNA-seq on a small number of asymptomatic WWWDs and closely-related redhead ducks, we found substantial differences in the expression of immune genes, including many genes associated with the RIG-I pathway, a critical component of innate immune response to TB (41), and the human tuberculosis KEGG pathway. One TB pathway gene, TLR2, is down-regulated in asymptomatic WWWDs relative to redhead ducks. TLR2 plays a critical role in the innate immune response. In mice, exposure to *M. tuberculosis* downregulates TLR2 and, in turn, interferes with the induction of the type I interferon response and cross processing by MHC Class I (20, 42). *Mycobacterium tuberculosis* itself produces TLR2 agonists in mice. We speculate that *M. avium* may have a similar effect in WWWD, whereas redhead ducks may be resistant to this form of immune system evasion. Necropsies of WWWDs have revealed broad dissemination of the disease across tissues, a high concentration of mycobacteria and a lack of multinucleated giant cells within lesions, all suggesting a failure of cell mediated immunity (43). MHC Class I, its heterodimer B2M, and associated transport proteins TAP1 and TAP2 were all upregulated in asymptomatic WWWDs relative to redheads, but no MHC genes were differentially expressed between symptomatic and asymptomatic WWWDs. This suggests that WWWDs have persistently engaged a core component (MHC Class I) of self–non-self recognition and adaptive immune pathways. This may be due to ubiquitous TB exposure in captivity. In this scenario, asymptomatic birds are actually pre-symptomatic; they are infected with TB and are mounting an immune response that is not effective at staving off disease in the long-term. Alternatively, these expression patterns could suggest underlying immune dysregulation such as autoimmunity or immune overexpression (44, 45).

Our findings strongly suggest that the genetic health of the captive WWWD population in the US is a key factor in their susceptibility to *M. avium*, but perhaps not the only factor. Husbandry changes, such as breeding birds at lower density, may be effective in reducing exposure to shed virus. Until now, vaccine development and administration has been prohibitively expensive, and a previous vaccination effort in the UK was not successful at preventing disease in adult WWWDs that had presumably been exposed to TB prior to vaccination (46). With the advent of mRNA-based vaccines for TB (47), cost-effective vaccines may be within reach and may be successful if aimed at TB-naive hatchlings. We suggest that a combination of genetic improvement, improved husbandry and vaccination should be pursued to improve captive breeding success. These efforts should be coordinated internationally, with an emphasis on working within the native range of the species, and maximising positive conservation outcomes in the wild.

## MATERIALS & METHODS

### DNA samples

Our sampling design included captive, India-derived birds from SHBP, and wild birds from Indonesia. Captive birds were sampled as part of routine health checks or during necropsies performed by veterinary staff at SHBP. These samples represent two time periods that we label “historical” and “current.” The current captive sample comprised ten individuals (six males and four females) hatched between 2008 and 2014. The historical captive sample included nine birds that were blood-sampled at Sylvan Heights in 1994. Among the birds in the historical sample, those with known studbook numbers hatched between 1985 and 1990 at three locations in North America; SHBP, NC; Goodewood Game Bird Farm, AL; and St. Louis Zoo, MO; all were subsequently moved to SHBP. Three individuals in the historical sample could not be associated with their corresponding studbook number and consequently have limited historical information.

Feather samples were collected from 11 wild WWWDs in Lampung Province, Indonesia, primarily in Way Kambas National Park on the southern tip of the island of Sumatra, between June 1999 and February 2001 by XXX. Samples were exported from Indonesia with permission from the Directorate General of Forest Protection and Nature Conservation in the Department of Forestry, and imported to the US under CITES permit 01US042484/9.

### DNA extraction, sequencing and read mapping

The ten current captive samples included DNA extracted from blood, liver, lung, and spleen. Nine current WWWD samples were extracted in 2017, with a Qiagen DNeasy Blood & Tissue kit. Historical captive WWWD DNA samples were extracted from blood samples collected at SHBP; standard phenol-chloroform extractions were completed by XXX in 1995 and stored in a -80ºC freezer before being sent to XXXXXXXXXX in 2019. The 11 wild WWWD feather samples collected by XXX were sent to XXX in 2003; DNA extraction from feathers was completed using the Qiagen DNeasy Tissue and Blood kit with the addition of 30 µl of 100 mg/ml dithiothreitol to digest the feather keratin; these extracts were preserved in a -80ºC freezer until being sent to XXXXXXXXXX in 2018 for whole genome sequencing.

Due to project funding and sample availability, samples were sequenced in two batches. Fragment libraries for a first batch of seven current samples were prepared with an Illumina TruSeq PCR-free kit and sequenced at the University of Illinois Roy J. Carver Biotechnology Center on an Illumina HiSeq 4000 instrument. A second batch of 22 birds, including the remaining current, historical and wild samples, were also sequenced at the University of Illinois. These libraries were prepared with a Hyper Library construction kit from Kapa Biosystems and sequenced on a NovaSeq 6000.

For most of the population genomic analyses reported here, WWWD data were aligned to the highly contiguous tufted duck genome (*Aythya fuligula* GCF_009819795.1, contig N50=17.7Mb), but we also mapped the data to the short-read-based WWWD genome assembly (GCA_013398475.1, contig N50=91.2kb (14)). Reads were aligned to both reference genomes using *bwa mem* v0.7.17(default settings) (48). Alignment success was similarly high for both reference genomes (WWWD reference: historical=99%, current=99%, wild=94%; tufted reference: historical=99%, current=99%, wild=95%). We favoured the tufted duck-aligned dataset in most downstream analyses because of our interest in multi-copy immune gene diversity, the annotations for which are better in contiguous genomes with high Contig N50 (49, 50). Depending on the given analysis we used either called genotypes from *GATK* or genotype likelihoods inferred by *ANGSD* v0.911 (51) and using the *ANGSD-wrapper (52)*. We estimated genotype likelihoods in *ANGSD* with the following parameters: -doGLF 3 -GL 1 -doMaf 1 -SNP_pval 1e6 -doMajorMinor 1 -minMapQ 30 -minQ 20 (51). Called genotypes, based on depth-normalized aligned data, were obtained using *bcftools* (v 1.13) *mpileup* and *bcftools call -m -f GQ (53)*, and included invariant sites. VCF files were filtered using *vcftools* v0.1.15 to include only sites that were present in 70% of individuals, and excluded sites with minDP<5 and max-meanDP>50. Both approaches yielded comparable results in downstream population genetic analyses, such as estimates of inbreeding and population divergence (not shown).

### Reconstruction of historical demography

We used *PSMC* (Li & Durbin 2011) to compare the demographic histories of the sampled WWWD populations. Demographic estimates from *PSMC* are impacted by variation in coverage, with lower coverage samples missing variants. Our captive birds had lower depth than our wild birds, so for these analyses we used our two highest depth captive birds and downsampled all the wild birds to match. This procedure will tend to reduce estimates of *Ne*, so our *PSMC* findings provide relative estimates of effective population size and population trajectory. For this analysis, we aligned trimmed reads to the reference genome using *bowtie v*2.5.4 and *–very-sensitive* options, we used *Picard* v3.1.1 *MarkDuplicates* to remove PCR duplicate reads. We then called variants for *PSMC* using *samtools mpileup* (-Q 30 -q 30 -u -v -f) for the 15 longest scaffolds and converted the output into a fastq using *vcfutils vcf2fq* (-d 4 -D 80 -Q 30). Hilgers et al (54) note a common issue in which PSMC estimates unreasonably large effective population sizes in the most recent time interval under typical settings. Following their guidance we split the first atomic time interval (-N25 -t15 -r5 -p “2+2+25*2+4+6”) which generated interpretable and reasonable demographic trajectories. PSMC results were scaled using a generation time of three years and a per-generation mutation rate of 2.5 × 10^-9^ (55).

### mtDNA assembly & phylogenetic reconstruction

To assemble the mitochondrial genomes, we aligned short read data to an existing mtDNA assembly for WWWD (GenBank accession: MN356440) and imported the resulting reads into Geneious Prime v. 2023.2.1 (https://www.geneious.com). Given the presence of nuclear copies of mtDNA sequences in related duck species (Sorenson et al. 1996), we closely examined alignments for evidence of “numts” (Sorenson & Quinn, 1998). For the feather samples from the wild population (*n* = 11), there was little evidence of numts and substantial coverage of the mtDNA (>800x), allowing inference of complete, high confidence mtDNA sequences for all samples. Results were more variable for blood samples from the captive population (*n* = 19 including the sample used for the reference genome), with coverage ranging from 22x to over 500x. Divergent numt reads were evident in all samples, but represented a decreasing proportion of reads as total coverage increased. Thus, complete mtDNA sequences could be assembled for those blood samples that yielded relatively high coverage of the mtDNA; this included 1 historical sample and seven current samples with total coverage of 85x or more. To provide an estimate of the timing of divergence of mtDNA lineages, we augmented the WWWD data with available waterfowl mtDNA genomes (see Figure Sx) and completed a phylogenetic analysis in BEAST (v. 2.7.7(56)) using the 12 light-strand encoded mt protein-coding genes. We partitioned the data by codon position, employed model averaging and an optimized relaxed clock (Bouckaert & Drummond 2017; Bouckaert et al. 2019). Following Mitchell et al. (57) and using identical parameters values, we time-calibrated the phylogeny by setting a lognormal prior on the age of the dabbling duck lineage based on the middle-Miocene fossil *Anas soporata* (58).

### Genetic diversity, inbreeding and differentiation

Using called genotypes, we estimated individual autosomal heterozygosity in each population using *vcftools* (--het) (59). We estimated nucleotide diversity (π) for each of the three populations and pairwise FST among populations using 100kb windows in *Pixy v2*.*0*.*0*.*beta6* (60). These analyses were based on VCF files with invariant sites included.

We used *ANGSD* and ANGSD-wrapper (51, 52) to visualize genetic differentiation with PCA and estimate inbreeding from genotype likelihoods. We estimated individual inbreeding coefficients using the expectation maximization algorithm in the ngsF module(61). We used *PCangsd* to represent genetic data as principal components (17).

Finally, we estimated runs of homozygosity in the genome using *ROHan v1*.*0*.*1* (62) directly from individual alignment files. We restricted the analysis to tufted duck autosomes longer than 10Mb and used the following parameters: rohmu=1×10^-3^, and window size=500kb. The genomic inbreeding coefficient (F_ROH_) was calculated as ∑L_ROH_/L_autosomes_ where ∑L_ROH_ is the total length of all autosomal ROHs in an individual and Lautosomes is the total length of the autosomal genome used in the analysis (63–65).

For captive birds only, we tested for associations between lifespan and observed heterozygosity, individual inbreeding coefficients and F_ROH_ using a gaussian generalized linear model in the R package *glmmTMB* and model checks in *DHARMa* (66, 67). We compared two models, one which included a parameter for historical versus current population and one with the genetic measure alone. Models were compared using AIC. Models and all plotting were conducted in *R* (V 4.4.0)(68)

### Major Histocompatibility Complex

We characterized diversity in functionally relevant peptide binding regions (PBR) of MHC classes Iα and IIβ. Our ability to accurately assess heterozygosity and MHC copy number may have been influenced by different factors depending on the reference genome we used. The WWWD genome assembly is likely incomplete for MHC genes, whereas the tufted duck genome may be divergent in copy number, such that some copies may have insufficient sequencing depth for accurate SNP calling (Figure S7). Therefore, we used two approaches to characterize MHC diversity in WWWDs. We first re-annotated the tufted duck genome using previously published consensus sequences of exons 2-4 derived from Anatidae MHC Class I and II PBRs (i.e., a total of six exons) (50). We extracted putative matching sequences with e-values <1e10 using *tblastn v2*.*12 (69)*, then extracted the hit with the lowest e-value when hits overlapped. We used Geneious Prime 2022 to visualize these sequences in the genomic context and checked for premature stop codons. We considered sequential copies of exons 2-4 as a complete copy. Using this method, we estimated that the tufted duck has nine copies of Class I and three copies of Class II, whereas the WWWD genome assembly has two and one copies, respectively. Class I estimates differ from the tufted duck NCBI annotation, which includes three genes, as many annotations contain multiple PBRs. Secondly, to ensure we did not underestimate diversity, we extracted a representative gene (all exons and introns) from the tufted duck genome and re-mapped all previous MHC-aligning depth-normalized reads to the single copies representing Class I and Class II, respectively. Using *samtools depth*, we assessed potential copy number by comparing the mean aligned read depth at the single copy MHC against the average depth of reads mapped to the tufted duck chromosome that contains MHC, chromosome 33. Copy number was assessed by dividing the average depth of MHC reads aligned to each MHC class and dividing by the average number of reads for chromosome 33.

Using this alignment, we called MHC SNPs for each class separately using *GATK HaplotypeCaller* v4.2.0(70, 71). Because multiple MHC copies were aligned to a single pseudoreference we doubled the copy number and used this number with the *--ploidy* flag. Then we filtered the VCF with the following expression: “QD < 2.0 || FS > 60.0 || MQ < 40.0 || MQRankSum < -12.5 || ReadPosRankSum < -8.0” (72). For this dataset, we estimated both nucleotide and amino acid diversity in PBR exons using custom code in R. For data aligned to the genome, we calculated nucleotide diversity using *pixy*, and calculated individual heterozygosity directly from genotypes in PBR exons.

### RNA samples

We sampled blood from eight WWWDs in 2022, and seven redhead ducks (*Aythya americana*) to provide a comparison to a closely related species that is less susceptible to avian TB. During the sampling period in 2022, two WWWDs were symptomatic for mycobacteriosis, whereas none of the redhead ducks were symptomatic. We further included two opportunistic samples from 2019: one redhead blood sample with mycobacteriosis, and a paired sample without (total redhead sample size=9). All instances of mycobacteriosis were fatal, and one WWWD was sampled approximately two hours post-mortem. This individual’s death was unexpected as it did not show outward symptoms of mycobacteriosis, but necropsy revealed typical mycobacterial lesions on its organs.

Birds were sampled at Sylvan Heights Bird Park using a 23 gauge needle and syringe to draw blood from the brachial vein. Whole blood was immediately placed in RNAlater®, and stored on ice until it was placed in a -80ºC freezer. All animal procedures were reviewed and approved by XXXXXXXXXX’s IACUC under Animal Use Protocol D351.

### RNA extraction, sequencing & read mapping

We extracted RNA from blood samples using TriZol and Qiagen RNeasy Kit, and all samples had RIN>7. Libraries were prepared with TruSeq® Stranded mRNA Library Prep, and sequenced on NextSeq 2000 P2 Reagents (200 cycles).

We generated 28.4-42.2 million paired-end reads per sample, retaining 89-97% of reads after trimming. We mapped RNAseq reads to the tufted duck genome (*Aythya fuligula*, GCF_009819795.1) using *STAR* v2.7.11b after trimming with *fastp* v0.23.2 (73, 74). Of these reads, 84-89% were uniquely mapped to the tufted duck genome (Figure S9). Reads mapping to each gene were counted using the *geneCounts* function in *STAR*.

### Gene expression

We filtered counts by removing genes with counts of ≤10 in at least six individuals. We identified three “potentially influential” individuals using Principal Component Analysis (using *PCAtools* (75)); these individuals had the smallest library sizes (Figure S9-S10). However, two of these three individuals represented two-thirds our disease sample, and so we used a conservative approach to remove the outlier effect from our list of differentially expressed genes (see below) We explored differences in normalized gene expression using a negative binomial generalized linear model framework implemented in *DESeq2* v1.44.0. First, we compared gene expression between asymptomatic redhead and white-winged ducks (gene expression ∼ species), using seven redhead samples (excluding one outlier individual) and six WWWD samples. Then, we compared diseased vs asymptomatic birds in each species separately (gene expression ∼ disease status), and compared the disease-related expression patterns between species. To identify differentially expressed genes (DEGs), we applied the Wald Test and considered genes to be significant with an FDR-corrected p-value (q) < 0.05. Due to the small sample sizes of diseased individuals, we removed significant genes that appeared to be affected by the outliers, rather than disease status, using custom code. We refer to the FDR-corrected p-value of genes that passed this additional filter as q*.

Susceptibility in WWWD ducks is likely linked to immune gene regulation or function, and thus we focused on immune DEGs. We extracted immune-related genes using a previously published list of immune GO terms (76), using the tufted duck GO annotations available on Genbank. We supplemented the immune gene list by including previously annotated MHC genes and searching the tufted duck reference annotations for genes relating to B-cells, T-cells, TLRs, immunoglobulin, interleukins, and interferons. We tested whether our candidate gene list was enriched in DEGs using the hypergeometric test function *phypher()* in R. To compare gene expression in the difficult multi-copy MHC, we summed counts from MHC Class Iα and MHC Class IIβ paralogs into a single “gene” to include in the analysis, but most reads mapped to a single copy in the tufted duck genome so this did not change the signal.

Then, we explored enrichment of GO Biological Processes and KEGG pathways using *clusterProfiler* V4.12.6 (77). We considered the full DEG list (q<0.05) as the foreground and the whole blood transcriptome as the background. For KEGG enrichment, we converted the NCBI gene-ids to functional ortholog ids (“ko”) using the *KEGGREST* v1.44.1 package (78), and used *enrichKEGG()* to identify enriched pathways. We excluded “significant” enrichments that included ≤ two genes.

## Data Availability Statement

Raw DNA sequence data are available under PRJNA1258233 (Table S1), raw gene expression data is available under PRJNA1285963 (Table S4). All data and code are provided in Dryad (https://doi.org/10.5061/dryad.crjdfn3ht).

