## Supplementary Figure for "Tuberculosis susceptibility and inbreeding depression hinder *ex-situ* conservation in a critically endangered rainforest bird"

Inbreeding depression and tuberculosis susceptibility in the critically endangered white-winged wood duck *Asarcornis scutulata*

Peri E. Bolton*, Dustin J. Foote*, Nancy Drilling, Susan B. McRae, Kim Cook, Michael D. Sorenson, Christopher N. Balakrishnan

**Supplementary Information and Figures**

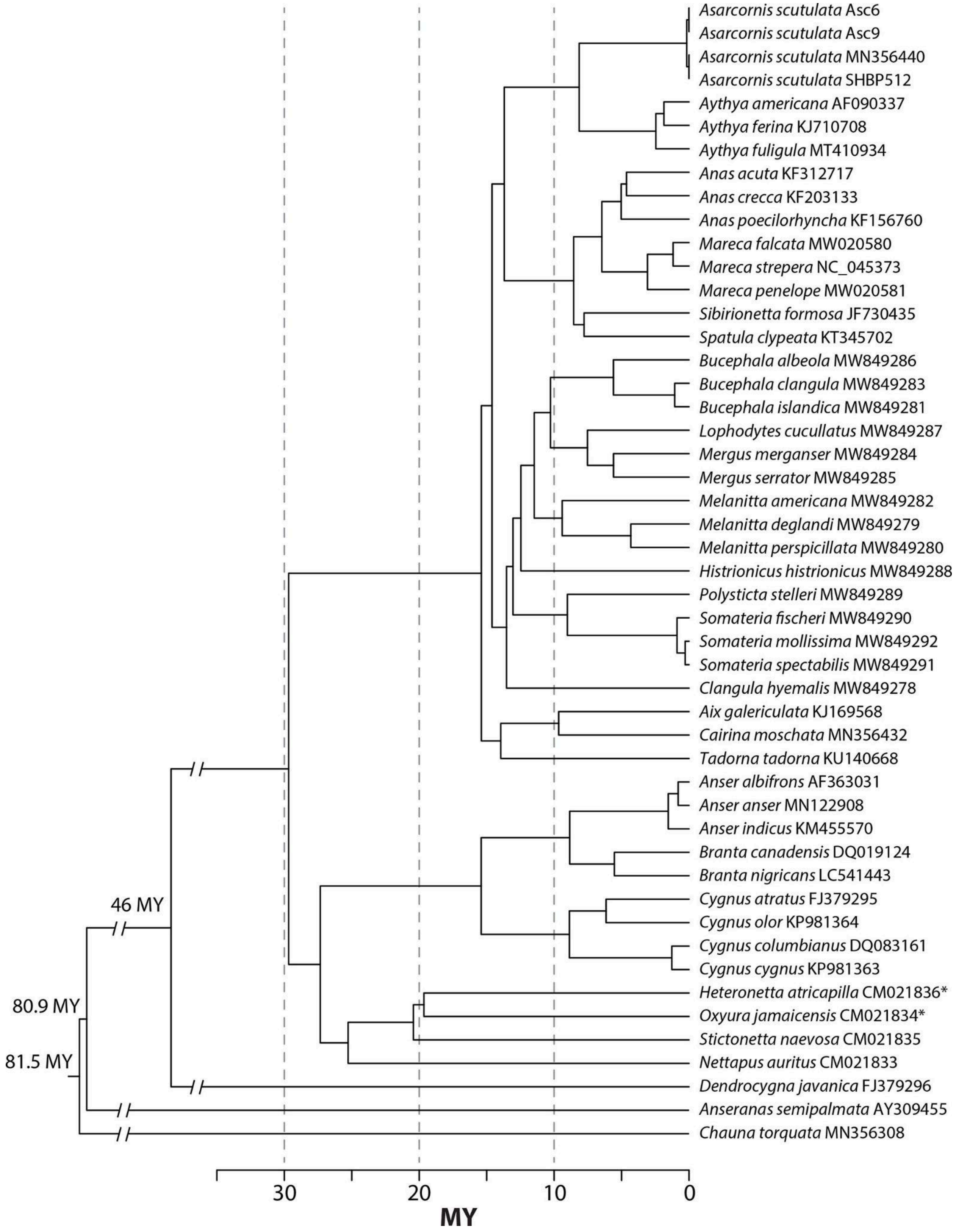


**Figure S1:** Time calibrated mtDNA phylogeny of ducks including Indian-derived and Indonesian White-winged wood ducks (*Asarcornis scutulata*, top).

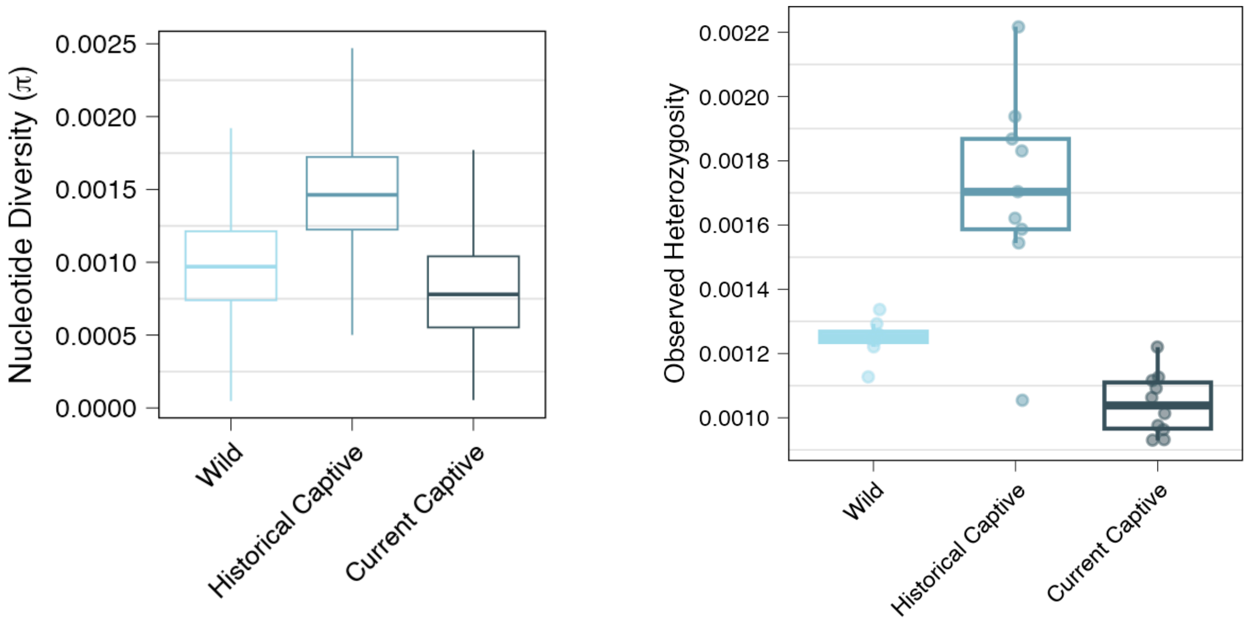


**Figure S2: Historical captive birds contain more diversity than wild birds in Indonesia and current captive birds.**
Left Panel: Nucleotide diversity calculated in 100kb windows across the genome.
Right Panel: Individual heterozygosity calculated using variant and invariant sites, where each point is an individual.

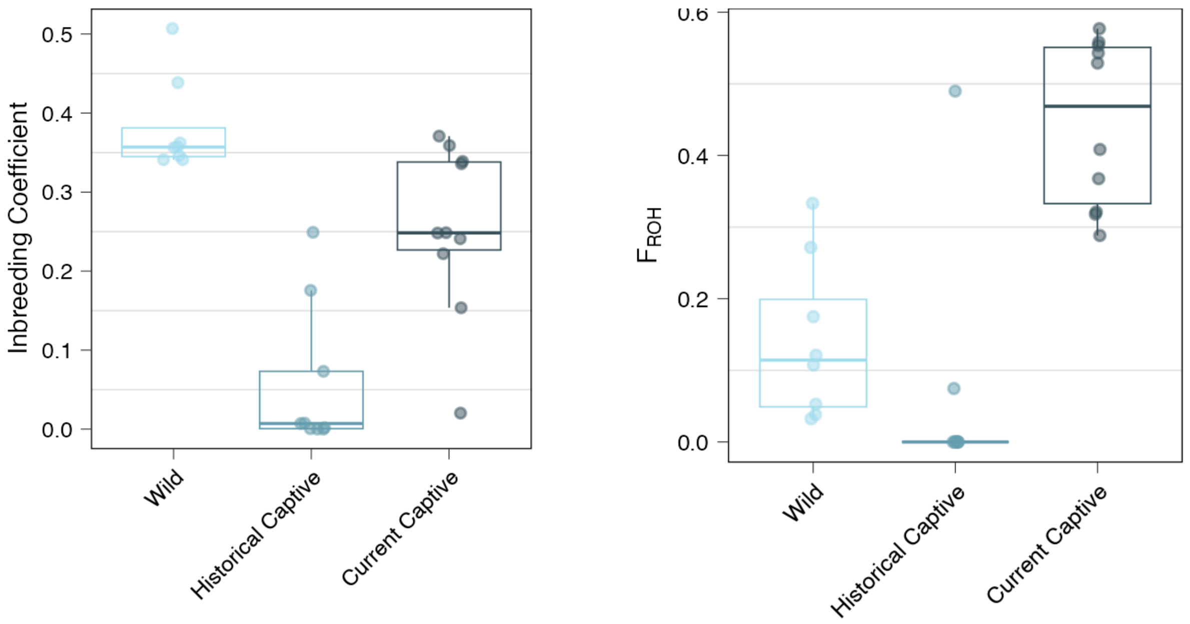


**Figure S3: Inbreeding coefficients from genome-wide heterozygosity using ngsF (left) and fraction of genome in ROHs (right).**
Each point represents the inbreeding coefficient of each individual in the sampled population.

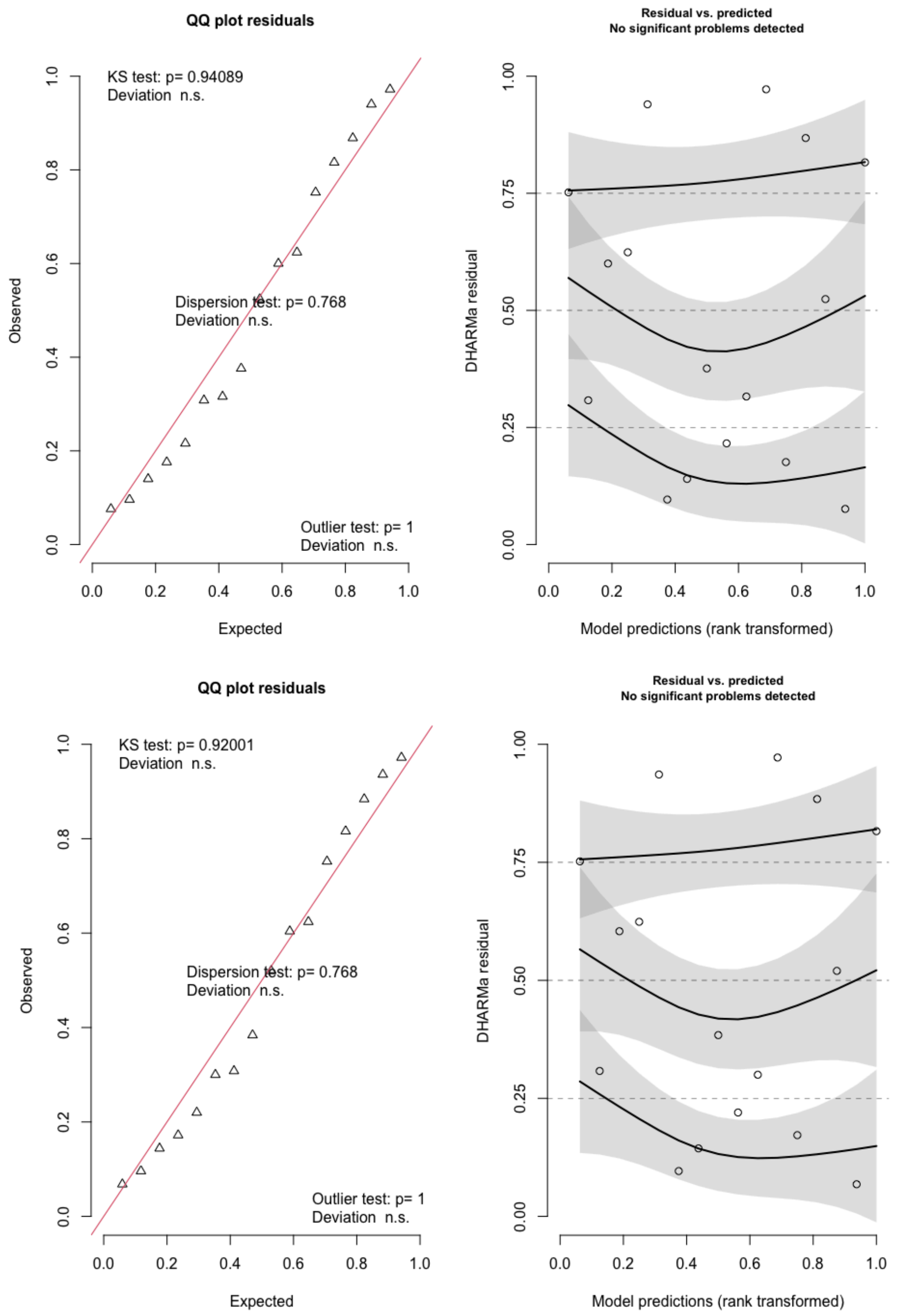


**Figure S4: Model assumptions checks from DHARMa indicating no significant departures from normality of residuals comparing inbreeding coefficient and lifespan. See also Table S2.**

Top model: inbreeding ~ population + age

Bottom model: inbreeding ~ age

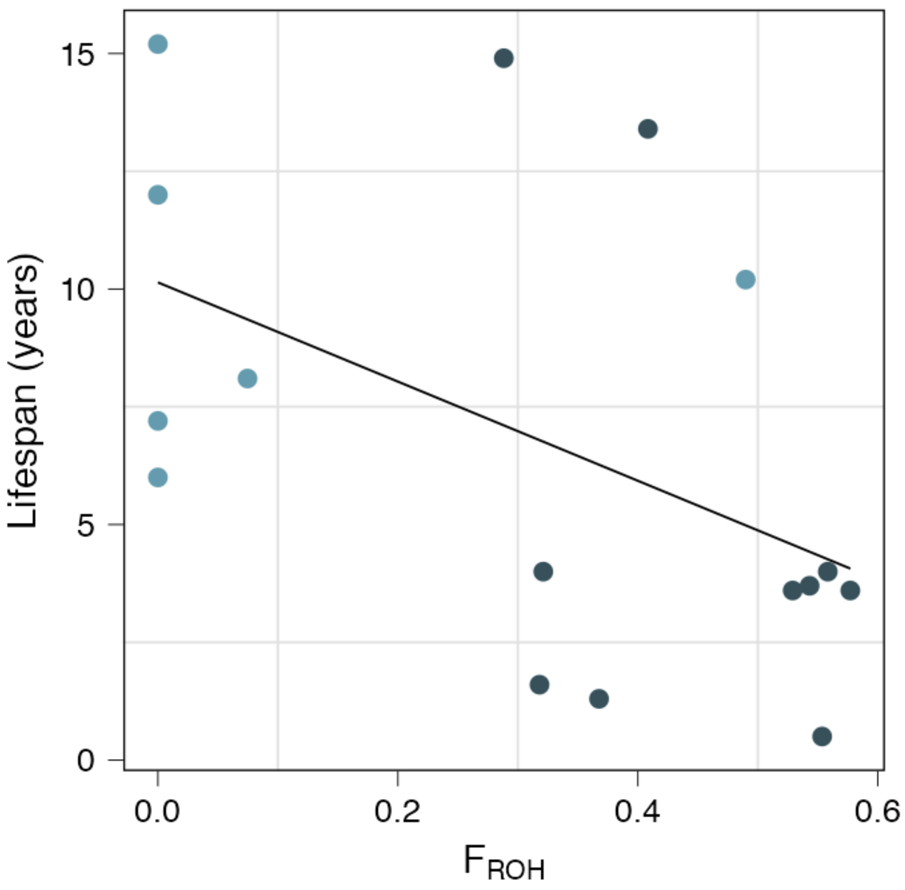


**Figure S5: Weak negative relationship between $F_{ROH}$ inbreeding coefficient and lifespan.**

Fitted line is from model age ~ $F_{ROH}$ ($r = -0.5$, $p(F_{ROH}) = 0.025$). Table S2 includes models outputs, and is indistinguishable by AIC from the nonsignificant model that includes a population variable. (age ~ pop + $F_{ROH}$: $p(F_{ROH}) = 0.45$).

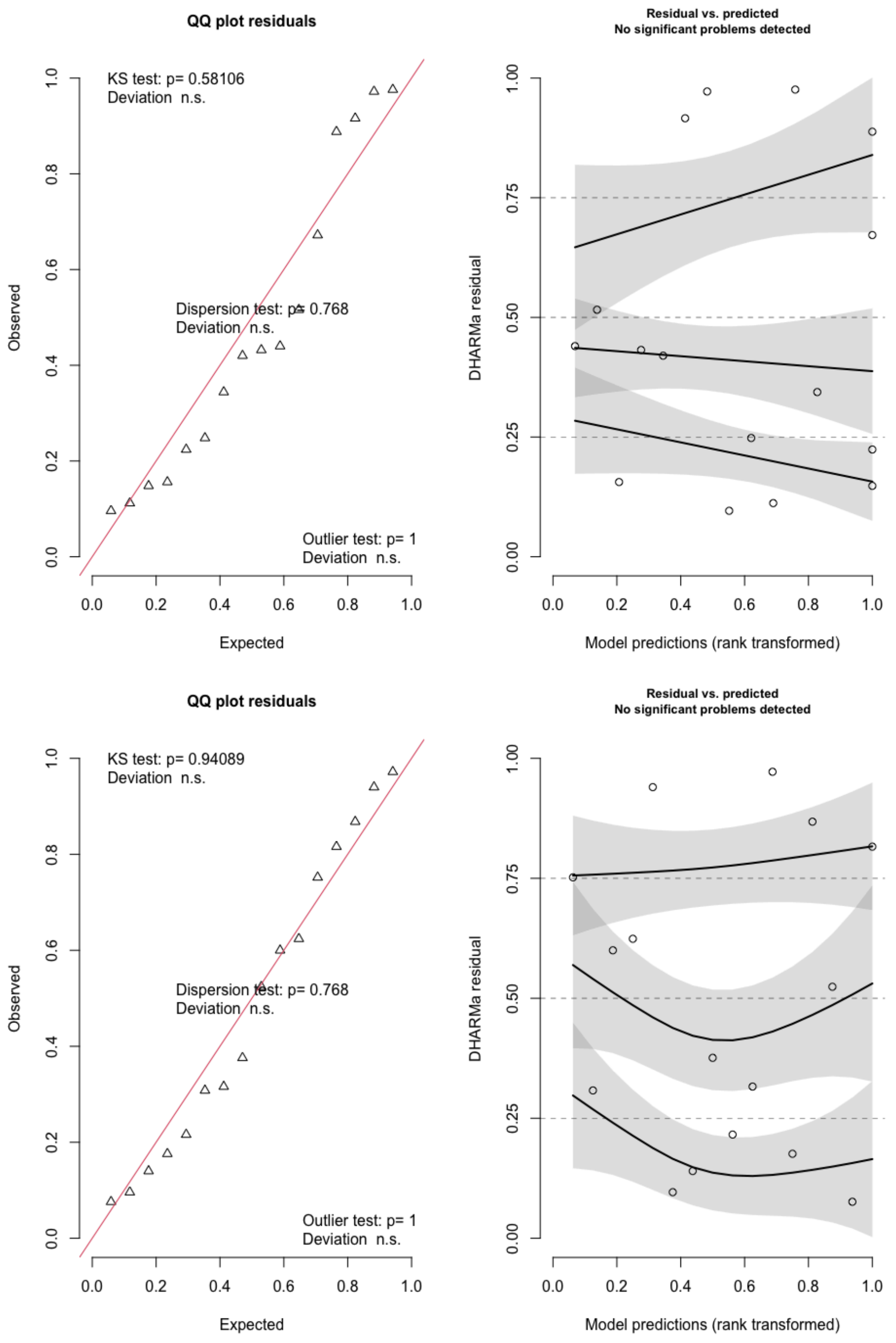


**Figure S6: Model assumptions checks from DHARMa indicating no significant departures from normality of residuals comparing $F_{ROH}$ and lifespan. See also Table S2.**

Top model: $F_{ROH}$ ~ age

Bottom model: $F_{ROH}$ ~ population + age

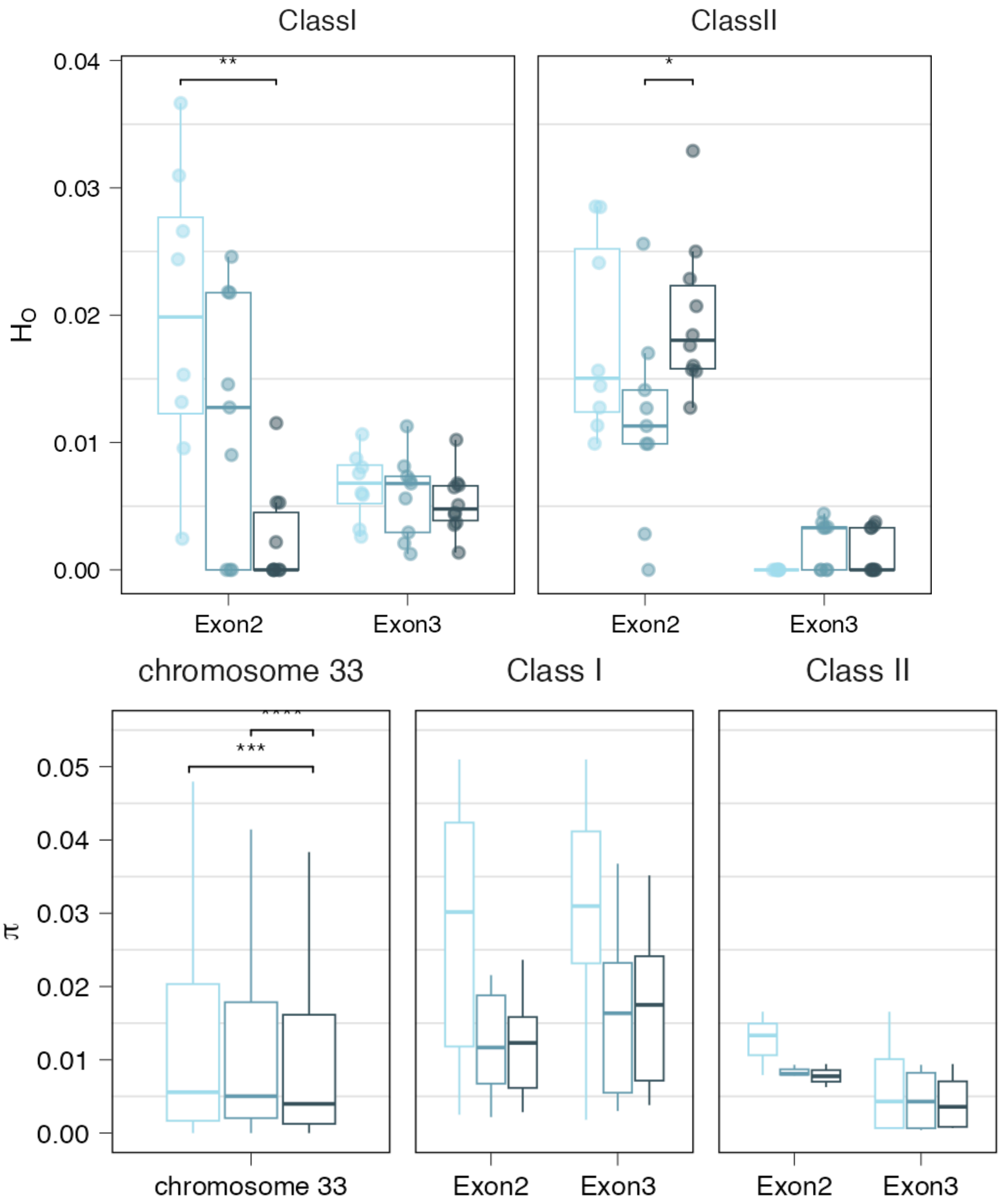


**Figure S7: heterozygosity (top) and nucleotide diversity (bottom) of MHC Peptide binding sequences when aligned to the tufted duck genome.** Colours (left to right/light to dark) are wild, historical captive, and current captive. We estimated heterozygosity directly from called genotypes for each individual as the number of heterozygous sites among variant and invariant sites in the exons. We calculated nucleotide diversity (π) in 1kb windows using pixy. Plots indicate significant wilcoxon tests adjusted for multiple testing with Holm method.

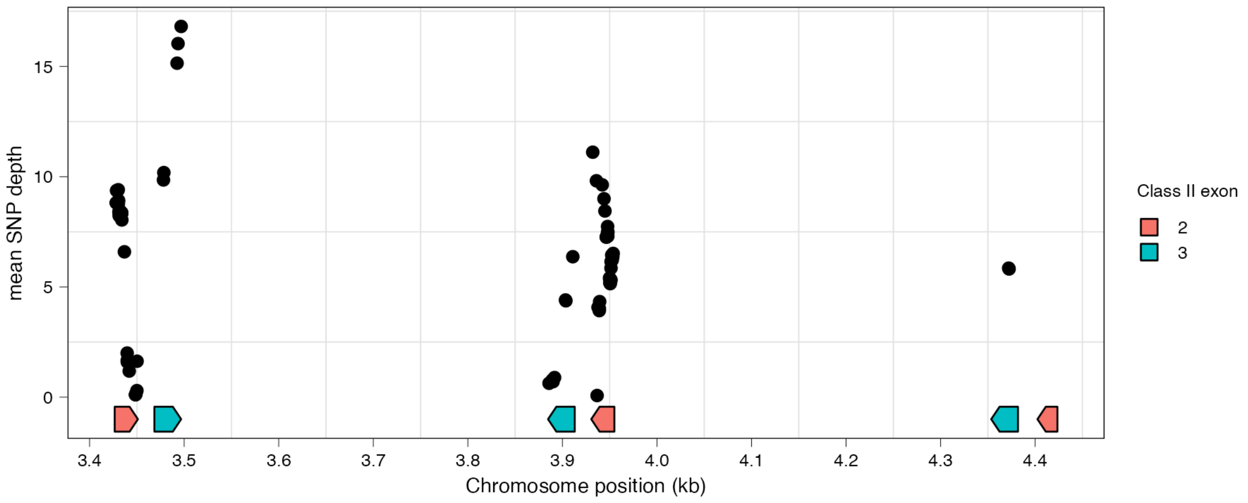


**Figure S8: SNP depth and density is inconsistent across MHC class II copies in the tufted duck genome.**
Figure is annotated with positions of class II exons, where each dot represents a SNP.

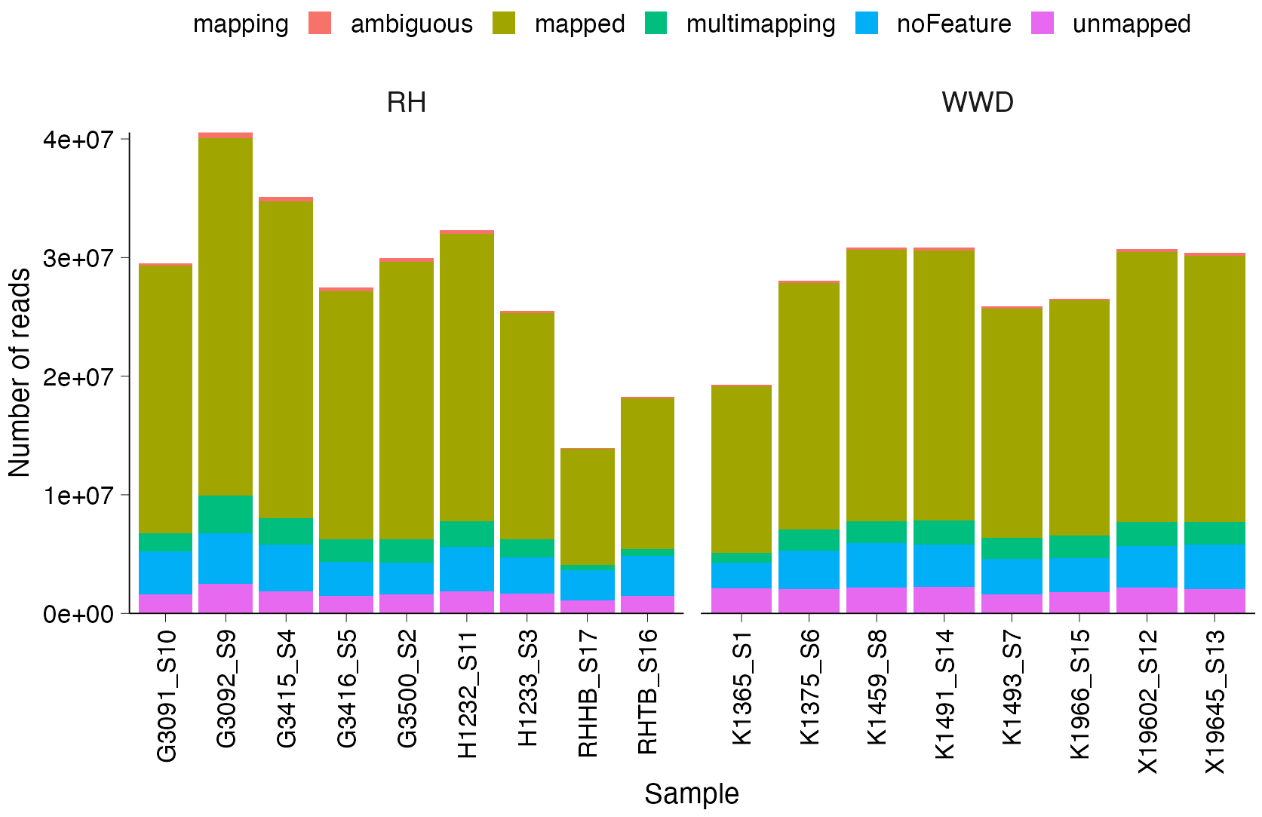


**Figure S9: Number of whole-blood RNAseq reads mapped to the tufted duck genome.**

Each sample is an individual redhead or white-winged duck, and includes the number of reads mapped to gene features in the *A. fuligula* genome (dark mustard colour). Other colours represent mapping flags and number of reads that were not included in downstream analyses.

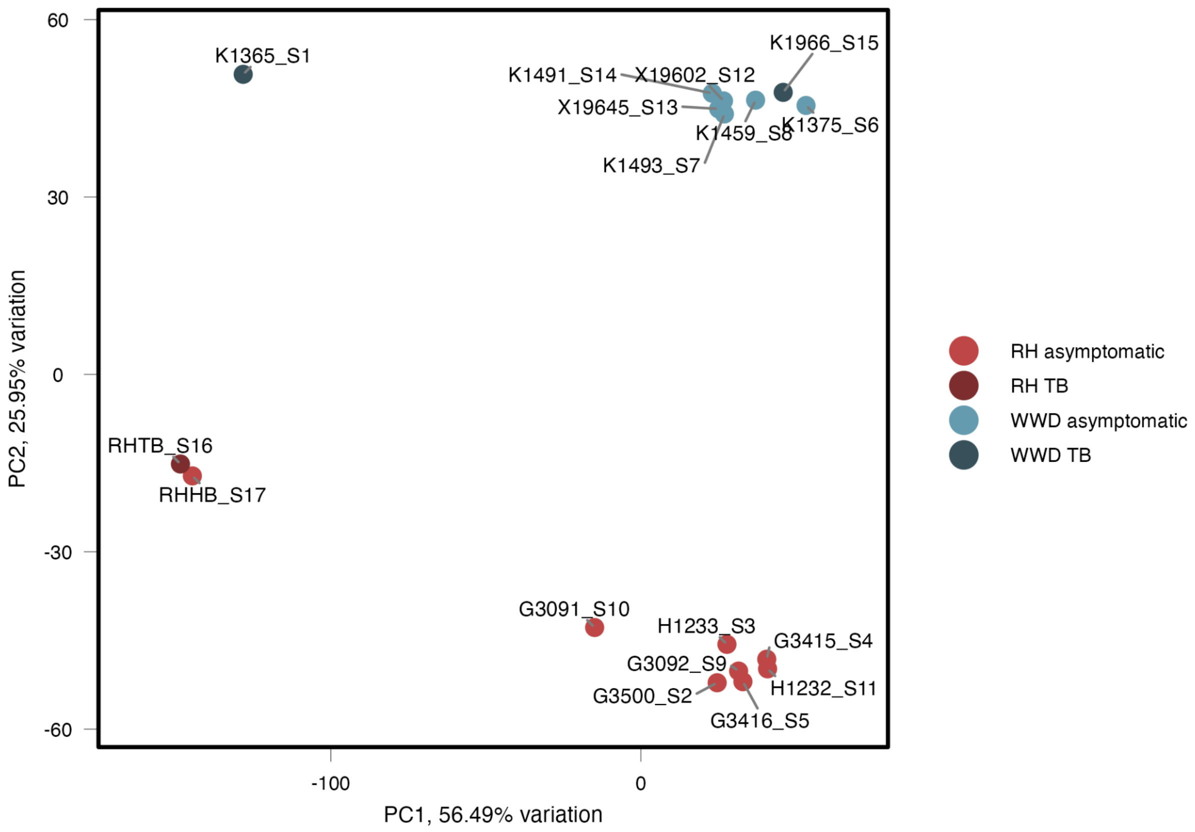


**Figure S10: PCA plot of all RNAseq samples after minimum count filtering.**
Each point is an individual's library, where colour represents species and point darkness indicates individuals that were symptomatic for mycobacteriosis (TB).

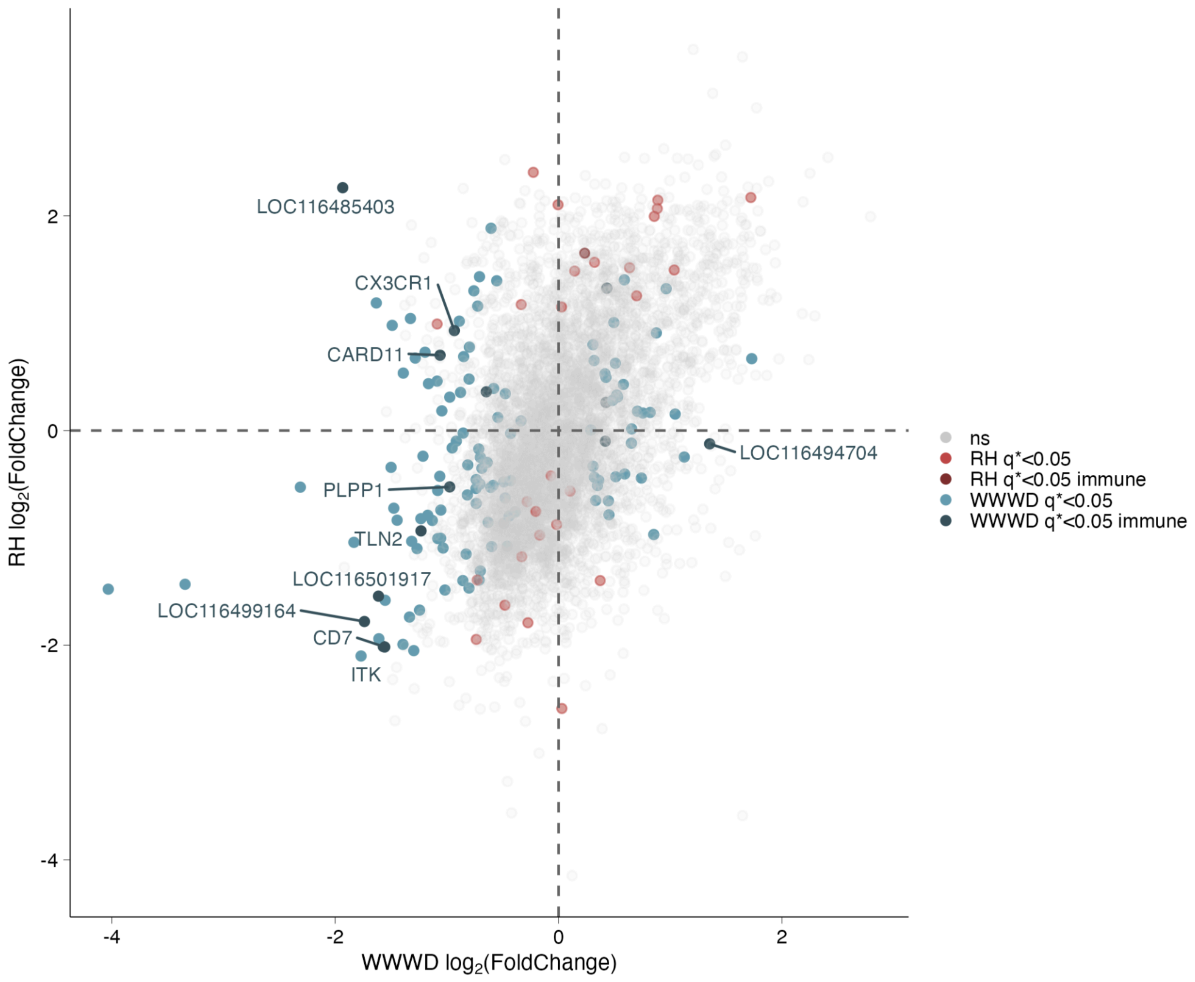


**Figure S11: Comparing disease expression profiles between redhead and WWWDs.**

The X-axis shows the direction and magnitude of expression of genes in symptomatic WWWDs, where positive values indicate upregulated in symptomatic individuals compared to asymptomatic individuals. Y-axis shows direction and magnitude of expression of genes in symptomatic redhead (RH) ducks. Genes that lie in bottom left and top right of the plot show similar expression profiles between species, while genes occupying the other quadrants show genes that are expressed differently in sick birds of these species. Coloured points are genes that were significant in each species' comparison and darker coloured points indicate immune system genes.
